# Preliminary evidence that maternal immune activation specifically increases diagonal domain volume in the rat brain during early postnatal development

**DOI:** 10.1101/432450

**Authors:** Tobias C. Wood, Michelle E. Edye, Michael K. Harte, Joanna C. Neill, Eric P. Prinssen, Anthony C. Vernon

## Abstract

Maternal immune activation (MIA) is consistently associated with elevated risk for multiple psychiatric disorders in the affected offspring. Related to this, an important goal of our work is to explore the impact of MIA effects across the lifespan. In this context, we recently reported the effects of poly (I:C)-induced MIA at gestational day (GD)15, immediately prior to birth, at GD21 and again at post-natal day (PD)21, providing a systematic assessment of plasma IL-6, body temperature and weight alterations in pregnant rats following poly (I:C) exposure and preliminary evidence for gross morphological changes and microglial neuropathology in both male and female offspring at GD21 and PD21. Here, we sought to complement and extend these data by characterising in more detail the meso-scale impact of gestational poly (I:C) exposure at GD15 on the neuroanatomy of the juvenile (PD21) rat brain using high-resolution, *ex vivo* anatomical magnetic resonance imaging (MRI) in combination with atlas-based segmentation. Our preliminary data suggest subtle neuroanatomical effects of gestational poly (I:C) exposure (n=10) relative to saline controls (n=10) at this time-point. Specifically, we report here preliminary evidence for a significant increase in the relative volume of the diagonal domain in poly (I:C) offspring (*p*<0.01; *q*<0.1), particularly in female offspring. This occurred in the absence of any microstructural alterations as detectable using diffusion tensor imaging (DTI). Longitudinal *in vivo* studies, informed by the effect sizes from this dataset are now required to establish both the functional relevance and cellular mechanisms of the apparent DD volume increase.

## 1. Introduction

Epidemiological studies report associations between exposure to maternal immune activation (MIA) and increased risk for psychiatric illnesses, including schizophrenia (SZ) and autism spectrum disorder (ASD) in the affected offspring [1, 2]. Based on these findings, reverse-translational animal models of MIA were developed, particularly (but not exclusively) using the viral mimetic polyriboinosinic-polyribocytidylic acid (poly I:C) [3, 4]. Administration of poly (I:C) on specific gestational days (GD; range 9–17) induces a discrete inflammatory response in the dam and elicits behavioural and neurochemical abnormalities in the offspring that are of relevance for the aforementioned psychiatric illnesses, providing both causation for the epidemiological data and important mechanistic insights [3, 5–7]. On the other hand, differences with regard to the strain of rodent, dose and route of administration of poly (I:C) used to induce MIA are present in the literature [5, 8–11]. These methodological differences likely explain the variance in published data with regard to behavioural and *post-mortem* brain phenotypes in rodent MIA models [8, 9, 11, 12]. Reporting guidelines and painstaking methodological work to establish the sources of variation in rodent MIA models, such as the caging system used to house the animals, are critical steps forward to address this issue [8, 10]. There is also a clear unmet need for an early outcome measure with which MIA-exposed offspring that will develop a robust behavioural phenotype may be identified and further enhance reproducibility across laboratories [12].

To address this gap, small animal magnetic resonance imaging (MRI) is one experimental approach, which offers several advantages. First, neuroanatomical phenotypes defined using high resolution MRI in combination with advanced image processing techniques are quite robust [13, 14]. Second, MRI provides whole brain coverage, eliminating the need for *a priori* hypotheses concerning implicated brain regions [15]. Third, MR imaging operates at mesoscopic resolution, which is technically translatable to human studies [16]. This is relevant in light of recent work examining the influence of maternal cytokine levels on brain structure and function in human offspring, in relation to the development of psychopathology [17, 18]. Fourth, non-invasive MR imaging in rodent models may be directly linked with behavioural assays and invasive *post-mortem* follow-up, to establish the neural correlates of behaviour and the underlying mechanism(s) [19]. The utility of this approach is exemplified by a number of landmark studies demonstrating that induction of MIA in rodents using a range of protocols is associated with consistent sex-specific deviations from the normative developmental trajectory of brain structure, function and neurochemistry [20–26]. As yet however, a reproducible non-invasive MRI biomarker measured in early life that is predictive of later dysfunction in MIA-exposed adult offspring has yet to emerge.

Related to this, an important goal of our work is to explore the developmental trajectory of MIA effects across the lifespan [27]. We have recently begun this process by exploring the effects of MIA prior to birth, at gestational day (GD)21 and just prior to weaning, at post-natal day PD21 [27]. This initial study (see also commentary by Roderick and Kentner, 2019 [28]) provided a robust and systematic assessment of plasma IL-6, body temperature and weight alterations in pregnant rats following poly (I:C) exposure and preliminary evidence for gross morphological changes (e.g. brain to body weight ratio, brain weight) and neuropathology (Iba1+ microglia number and morphology) in male and female offspring at GD21 and PD21. The relevance of these changes is currently being confirmed by behavioural assessment [27]. Emerging evidence however, shows behavioural changes of relevance to psychiatric disorders in GD15 poly (I:C) exposed offspring, including an increase in ultra-sonic vocalisation in early life (PD6) and a deficit in sustained attention in adulthood (PD125) [29]. Here, we sought to complement and extend these initial results by characterising the impact of MIA on brain volume and microstructure using high-resolution, *ex vivo* anatomical magnetic resonance imaging (MRI) specifically at PD21. The rationale for this approach is two-fold. First it will provide a detailed, brain-wide assessment of early neuroanatomical differences between control and MIA-exposed offspring, using the protocol reported in [27]. Second, it will provide preliminary data (i.e. effect sizes) to inform the design of future longitudinal *in vivo* MRI studies, which are necessary to establish if early MRI-detectable brain changes have any functional relevance.

## 2. Methods

### 2.1 Animals

Animals used in the present study were taken from a large cohort of control and poly (I:C)-exposed Wistar rats generated at the University of Manchester. Full details of rat strain, sex and housing conditions are reported in detail elsewhere [27]. All procedures in this study were carried out in accordance with the UK Home Office, Animals (Scientific Procedures) Act 1986 and EU Directive 2010/63/EU. The University of Manchester Animal Welfare and Ethical Review Body (AWERB) approved all experimental protocols used in this study.

### 2.2 Maternal immune activation (MIA) and allocation of offspring for ex vivo MRI phenotyping

Induction of MIA was performed at the University of Manchester as described in detail elsewhere [27]. In brief, female Wistar rats were mated at 3 months of age and GD1 confirmed by the appearance of a vaginal plug. Several studies provide evidence that poly (I:C) treatment at GD15 in rats induces a maternal inflammatory response with development of relevant behavioural phenotype(s) in the affected offspring [24, 26, 30, 31]. Pregnant Wistar rats (293.0–428.7 g) therefore received a single intraperitoneal (i.p.) injection of either poly (I:C) (n=8; P9582, potassium salt, Sigma-Aldrich; Gillingham, Dorset, United Kingdom) at a dose of 10 mg/kg or 0.9% non-pyrogenic sterile saline as a control (n=8) on GD15. On PD1, pups were sex typed based on anogenital distance and then culled to litters of n=4 males and n=4 females. At PD21 all pups were sacrificed and n=1 male and n=1 female offspring selected from 5 (out of the available 8) control and poly (I:C) litters, giving a total of n=10 (5 male, 5 female) offspring per experimental arm (control vs. poly (I:C)). In the absence of data concerning effects of MIA on brain volume in Wistar rats at PD21 and how this may vary by sex, a formal estimation of sample size by power calculation was not possible. Rather, this study was designed pragmatically to generate this data using the minimum number of animals, in line with UK and EU guidelines (see www.nc3rs.org.uk) and taking into account available MRI scanner time and financial resources. We therefore selected a group size of n=10, comprised of 5 males and 5 females, such that both experimental groups were matched for sex. This group size (n=10) is comparable with prior *in vivo* neuroimaging studies in rodent MIA models [20, 24–26, 32]. As such, the results presented herein are preliminary given the limited sample size and should be interpreted accordingly (*see also discussion*).

### 2.3 Enzyme-linked immunosorbent assay

To confirm successful MIA in the dams, IL-6 concentrations in maternal blood plasma (3 hours post-injection) were determined using a rat-specific ELISA DuoSet (R&D Systems, Abingdon, UK) as reported elsewhere [27]. In brief, absorbances were measured using a plate reader (MRX, Dynatech, UK) at room temperature and results were calculated from the standard curve using Prism software (v6.0, GraphPad, La Jolla, CA, USA). We present here only the IL-6 values (pg/ml) for the specific dams from which offspring were selected for inclusion in the MRI study.

### 2.4 Tissue preparation for ex vivo MRI

At PD21, offspring were culled by cardiac perfusion (0.9% saline followed by 4% paraformaldehyde) under terminal anaesthesia (sodium pentobarbital, 60 mg/kg i.p) and the brain tissue prepared for *ex vivo* MRI as described elsewhere [16]. In brief, fixed brain tissues were kept intact in the cranium and post-fixed for 24 hours in 4% PFA. Samples were then placed in 0.01M phosphate buffer containing 0.05% (w/v) sodium azide to allow tissue re-hydration prior to MR imaging. Samples were then shipped to KCL and stored at 4°C in this solution for 4 weeks prior to MR imaging.

### 2.5 MR image acquisition

A 7T horizontal small bore magnet (Agilent Technologies Inc. Santa Clara, USA) and a quadrature volume radiofrequency coil (39 mm internal diameter, Rapid Biomedical GmbH, GER) were used for all MRI acquisitions. Fixed brain samples were placed securely one at a time in a custom-made MR-compatible holder and immersed in proton-free susceptibility matching fluid (FluorinertTM FC-70; Sigma-Aldrich, UK). Samples were scanned in a random order, with the KCL operator (ACV) blinded to treatment group (CON or POL) by numerical coding of samples. Scanning was interspersed with phantoms to ensure consistent operation of the scanner. Two sets of MR images were acquired: a 3D Fast-Spin Echo (FSE) for structural analysis and a DTI protocol for microstructural analysis. The T2-weighted 3D FSE image had the following parameters: TE/TR=60/2000, echo train length = 8, matrix size = 192×128×192 and field of view (FOV) = 28.8×19.2×28.8 mm, yielding isotropic voxels of 150 *μ*m^3^. Total scan-time was 1hr 44min. The DTI scans were acquired using a four shot Echo Planar Imaging (EPI) sequence with TE/TR=35/4000ms, matrix size = 128×96 and FOV = 25.6×19.2mm, with an in-plane resolution of 200 μm across 50, 0.5 mm-thick slices. A total of 30 non-collinear diffusion directions were acquired, with four *b*=0 images with a target *b* value of 2000 s/mm. Total scan time was 1hr 22min. Reversed phase-encoded direction DTI images were additionally acquired to correct for eddy current distortions (8min). Total scan duration for the entire protocol was 3hr 14min per brain.

### 2.6 MR image processing

After visual inspection of all MR images and elimination of those scans with artefacts, the final n values per group for statistical comparisons were: volume; CON, n=9 (4 male, 5 female) vs. POL, n=10 (5 male, 5 female); DTI: CON, n=10 (4 male, 4 female) vs. POL, n=10 (5 male, 5 female). The MR Images were converted to NIFTI format and processed using a combination of FSL [33], ANTs [34] and in-house C++ software utilizing the ITK library, available from https://github.com/spinicist/QUIT. The processing pipeline consisted of several steps, as described elsewhere [35]. The following operations were carried out in the native space of the acquired MR images. First, a Tukey filter was applied to the FSE images in k-space to remove high frequency noise and they were corrected for intensity inhomogeneity using the N4 algorithm [36]. Second, FSL *topup* and *eddy* were used to remove distortion and eddy current artefacts in the raw diffusion data [37, 38] using acquired DTI data with a reversed phase-encode direction (*see section 2.5*). The DTI parameter maps were then calculated using FSL *dtifit* and consisted of Fractional Anisotropy (FA) and Mean Diffusivity (MD) [35]. Third, a template image was constructed from the 3D FSE images of all subjects in the study using the *antsMultivariateTemplateConstruction2.sh* script with cross-correlation metric and SyN transform (n=19) [31]. This template was then registered to an atlas image of the PD18 Wistar rat brain, again using a cross-correlation metric and SyN transform [39]. Fourth, all subjects FSE images were non-linearly registered to the study template using the *antsRegistrationSyN.sh* script. Logarithmic Jacobian determinants were calculated from the inverse warp fields in standard space to estimate apparent volume change, and smoothed with a Gaussian filter at a full-width half-maximum (FHWM) of 200 *μ*m [40]. Fifth, each subject’s DTI image was registered to the same subject’s FSE image using a SyN transform to account for residual distortions and a Mutual Information metric to account for the different contrast. This transform was then concatenated with those from the FSE images to the templates to align them to the study template [35]. The DTI images in the template space were also smoothed with a Gaussian filter with full-width half-maximum of 200 μm. Sixth, a brain parenchyma mask was created from the atlas labels by excluding Cerebrospinal Fluid (CSF) regions. The inverse transforms from the atlas to the study template and from the study template to each subject were applied to calculate the brain and atlas based region of interest (ROI) volumes for each subject [35].

### 2.7 Statistical analysis

A subset of maternal plasma levels of IL-6, consisting of those values only from dams whose offspring were selected for MR imaging (n=5 CON dams vs. n=5 POL dams) were compared using unpaired t-test (2-tailed), with α=0.05, using Prism software (v6.0; GraphPad Inc. La Jolla, CA, USA). For the MR image analysis, group-level differences in volume, FA (unit less) and MD (mm^2^ s^−1^) were assessed using atlas-based segmentation (ABS) [20], taking advantage of a publicly available high resolution MRI atlas of the Wistar rat brain at PD18 that is parcellated into 26 regions of interest (ROI) [39]. We analysed relative volumes since total brain volume accounts for the majority of inter-animal variation in the volume of individual brain structures [13, 41]. Volume data for each atlas ROI was therefore expressed as a percentage of total brain volume [20]. FA and MD values were analysed without this correction. After image registration we successfully extracted values for volume, FA and MD for 23/26 ROIs (missing ROIs: Pineal gland, pituitary gland, optic pathway). Total brain volume was calculated from the summation of each individual atlas ROI volumes [20]. Group-level differences between CON and POL-exposed animals in relative volume, FA and MD were assessed across these 23 ROI using a 2 x 2 ANOVA, with MIA (CON or POL) and sex (male or female) as between and within group factor respectively, using Prism software (v6.0; GraphPad Inc. La Jolla, CA, USA) with α=0.05. The resulting *p*-values from the F-test contrasts (main effects of prenatal treatment, sex and prenatal treatment x sex interaction) were then corrected for multiple comparisons (to account for Type I errors across the 23 ROI) using the False Discovery Rate (FDR) procedure, with the threshold set at 10% (*q*<0.1). This threshold was chosen based on simulation studies showing that, at a 10% FDR, simulated volume losses/gains of 5–20% can be recovered with a true false positive rate of <0.2% [42]. Therefore in addition to *p*-values, we report *q*-values, which are FDR-adjusted *p*-values. Effect sizes were calculated using either partial η^2^ (ANOVA) or Cohen’s *d* (t-test) as appropriate. Due to the small sample size, particularly for sex, we do not present any voxel-wise, whole brain analysis.

## 3. Results

### 3.1 Maternal circulating IL-6 levels

A statistically significant increase in the circulating levels of IL-6 in maternal plasma samples could be observed 3 hours post-injection in POL-injected dams as compared to controls (*t*=3.62, *df*=14; *p*=0.007; Cohen’s *d*=2.3; **Figure 1**). There was however, a notable degree of variability in the plasma IL-6 levels within the POL-injected dams (range 480 to 2890 pg/ml).

**Figure 1.**
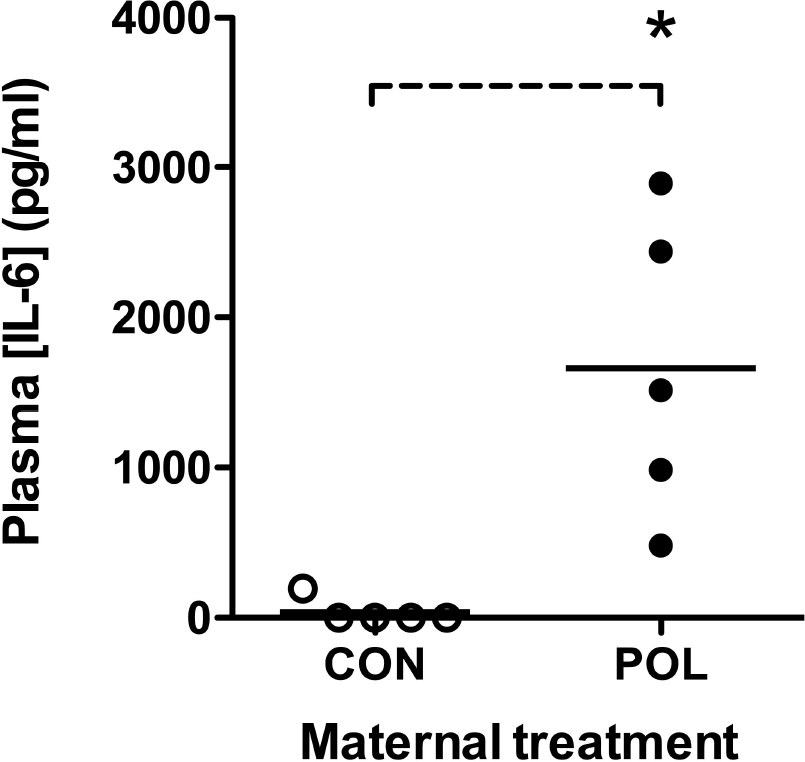
Significant increases in circulating maternal IL-6 levels 3 hours post-poly (I:C) (POL) injection (plasma samples from n=5 dams; 10 mg/kg i.p. administered on GD15) as compared to saline-injected dams (CON; plasma samples from n=5 dams; sterile saline, i.p.) determined using ELISA-based assay. Data shown are IL-6 levels in pg/ml, *p<0.05; 2-tailed student’s t-test.

### 3.2 Atlas based segmentation analysis of volumetric and DTI datasets

#### 3.2.1 Juvenile rat brain volume

Whole brain volumes were significantly affected as a function of sex (F(1,15)=5.89 *p*=0.03). There were no significant effects of either MIA, or MIA x sex interaction (**Table 1**). Main effects of sex were also found for the relative volumes of 3/23 (13%) atlas ROIs, including the bed nucleus of the stria terminalis (BNST), the cerebellum and the ventricular system, although these did not survive correction for multiple comparisons (**Table 1**). Nonetheless, these trend-level results are consistent with known sexually dimorphic regions in the rodent brain [43, 44]. A significant main effect of MIA was found for the relative volume of the diagonal domain (DD), with subtly increased volume in POL as compared to CON offspring (+4.4%; *p*<0.05, *q*<0.1; partial η^2^ = 0.44; **Table 1**; **Figure 2**). The relative volumes of the olfactory structures and preoptic area were also affected by MIA exposure, albeit only at trend level (*p*<0.05, *q*>0.1; Table **1**). Interactions between sex and MIA were found, again only at trend level, for the relative volumes of a further 3/23 (13%) of the atlas ROIs, including the DD, nucleus accumbens and anterior commissure (*p*<0.05, *q*>0.1; **Table 1)**.

**Table 1.** The results of a 2 x 2 ANOVA statistical analysis of whole and regional brain volumes at PD21 comparing CON and POL-exposed offspring. Only ROIs with *p*<0.05 (prior to multiple comparison correction) are shown. The *p*-values shown are derived from the F-tests of the main effects of either MIA or sex and MIA x sex interaction run across all 23 ROI. The *q*-values indicate statistical significance after FDR correction using the Benjamini-Hochberg procedure with a 10% threshold (*q*<0.1) to control for Type-I errors across the multiple ANOVA tests. Significant (*p*<0.05; *q*<0.01) F-tests are highlighted in red, whilst those in blue were significant at trend-level only (*p*<0.05; *q*<0.1).

| Region of interest | 2 x 2 ANOVA |  |  | Post-hoc test |
| --- | --- | --- | --- | --- |
|  | Within subjects |  | Between groups |  |
|  | Sex (*p, q*) | MIA x Sex interaction (*p, q*) | MIA (*p, q*) |  |
| Whole brain | *F*(1,15)=5.89; *p*<0.05 (*q*>0.1) | *F*(1,15)=3.32; *p*>0.05 (*q*>0.1) | *F*(1,15)=0.31; *p*>0.05 (*q*>0.1) | None |
| BNST | *F*(1,15)=4.97; *p*<0.05 (*q*>0.1) | *F*(1,15)=0.16; *p*>0.05 (*q*>0.1) | *F*(1,15)=0.01; *p*>0.05 (*q*>0.1) | None |
| Cerebellum | *F*(1,15)=4.40; *p*<0.05 (*q*>0.1) | *F*(1,15)=1.25; *p*>0.05 (*q*>0.1) | *F*(1,15)=0.44; *p*>0.05 (*q*>0.1) | None |
| Lateral ventricles | *F*(1,15)=4.34; *p*<0.05 (*q*>0.1) | *F*(1,15)=0.09; *p*>0.05 (*q*>0.1) | *F*(1,15)=0.01; *p*>0.05 (*q*>0.1) | None |
| Diagonal domain | *F*(1,15)=0.12; *p*>0.05 (*q*>0.1) | *F*(1,15)=5.91; *p*<0.05 (*q*>0.1) | *F*(1,15)=11.8; *p*<0.01 (*q*<0.1) | None |
| Olfactory structures | *F*(1,15)=0.13; *p*>0.05 (*q*>0.1) | *F*(1,15)=1.07; *p*>0.05 (*q*>0.1) | *F*(1,15)=6.11; *p*<0.05 (*q*>0.1) | None |
| Preoptic area | *F*(1,15)=0.02; *p*>0.05 (*q*>0.1) | *F*(1,15)=0.03; *p*>0.05 (*q*>0.1) | *F*(1,15)=4.77; *p*<0.05 (*q*>0.1) | None |
| Nucleus accumbens | *F*(1,15)=1.06; *p*>0.05 (*q*>0.1) | *F*(1,15)=5.49; *p*<0.05 (*q*>0.1) | *F*(1,15)=2.19; *p*>0.05 (*q*>0.1) | None |
| Anterior commissure | *F*(1,15)=0.41; *p*>0.05 (*q*>0.1) | *F*(1,15)=5.11; *p*<0.05 (*q*>0.1) | *F*(1,15)=0.10; *p*>0.05 (*q*>0.1) | None |

**Figure 2.**
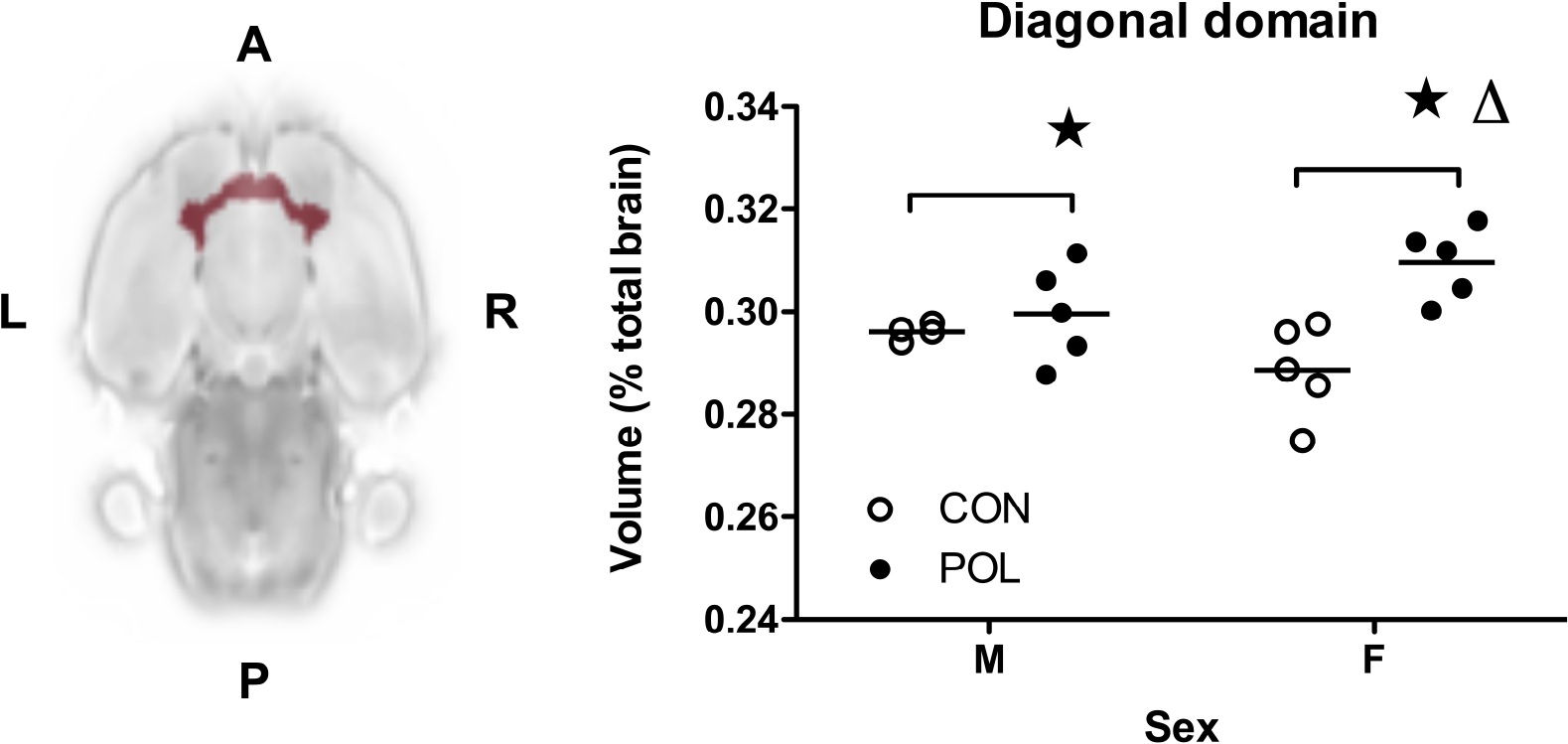
Prenatal exposure to POL at GD15 resulted in a significant main effect of MIA (*p*<0.01; *q*<0.1; partial η^2^ = 0.44) on the relative volume of the diagonal domain (DD) with an increase in POL-exposed offspring compared to controls at PD21. This increase is more apparent in the females POL offspring as compared to the males. Data shown are relative volumes, expressed as the percentage of total brain volume. ⋆*p*<0.01 main effect of MIA, corrected for multiple comparisons at *q*<0.1 using FDR procedure; Δ*p*<0.05, MIA x sex interaction at trend level significance (*p*<0.05; *q*>0.1). **A**, Anterior; **P**, posterior; **L**, left, **R**, right.

#### 3.2.1 Juvenile rat brain microstructure (FA, MD)

There were no statistically significant effects of sex, MIA or MIA x sex interaction on either FA or MD, across any of the 23 ROIs in the Wister PD18 MRI atlas (*all p>0.05; q>0.1; data not shown*).

## 4. Discussion

The main findings of the current study are as follows. Exposure of pregnant Wistar rat dams to poly (I:C) (POL) at GD15 has no effect on the total brain volume (in mm^3^) of juvenile offspring at PD21 relative to controls (CON). Irrespective of gestation treatment brain volume did vary as a function of sex, which is consistent with published data [43, 44]. When comparing the relative volumes of 23 brain regions between CON and POL offspring at PD21, we found only subtle neuroanatomical differences in the juvenile Wister rat brain.

Specifically, we provide preliminary evidence for a significant increase in the relative volume of the diagonal domain (DD) in POL offspring with a robust effect size (partial η^2^ = 0.44). This effect appeared to be driven primarily by female POL offspring, although this result did not survive FDR correction (MIA x sex interaction; *p*<0.05; *q*>0.1). Additional volumetric changes, albeit at trend-level only, were found for the relative volumes of the olfactory structures and pre-optic area. Finally, we found no significant differences, even at trend-level, for microstructural alterations as indexed by either FA or MD. Taken together, these data provide new information on the impact of MIA on the juvenile Wistar rat brain, complementing and extending our recently published work using this MIA protocol [27].

Before interpreting these findings, limitations of the current study should be noted. First, behavioural phenotyping of offspring generated using the MIA protocol used herein and in our prior study [27] is currently ongoing; as such the functional relevance of this MIA protocol still remains to be confirmed. Emerging evidence however, shows behavioural changes of relevance to psychiatric disorders in GD15 poly (I:C) exposed offspring, including an increase in ultra-sonic vocalisation in early life (PD6) and a deficit in sustained attention in adulthood (PD125) [29]. Nonetheless, we interpret our current results only within the narrow framework of further defining the impact of a maternal systemic poly (I:C) challenge of 10 mg/kg at GD15. Second, as already fully acknowledged (*see methods, section 2.2*) our sample size is small, precluding the full use of voxel-wise MR image analysis tools [20]. This could account for the limited number of significant findings surviving multiple comparisons correction, particularly with regard to MIA x sex interactions. Equally, we cannot exclude the possibility that our negative results, particularly where the DTI data are concerned, are not simply a reflection of the small sample size. As already stated, the group sizes were chosen pragmatically based on the absence of prior data regarding effect sizes, ethical use of the minimum number of animals, access to the MRI scanner and financial restraints. Our overall group size is also comparable to previously published MRI studies in MIA models, but we acknowledge this is likely underpowered to detect sex-specific effects [20, 24]. Taking these factors into account we refer to all findings as preliminary, pending replication in a larger cohort, which the data generated in the current study allows us to accurately calculate. For example, using the effect size for the main effect of MIA on relative DD volume (partial η^2^ = 0.44), a power calculation (F-test, repeated measures ANOVA, between factors) suggests that for a longitudinal imaging study with 2 groups (CON and POL) and 3 time-points, at least n=16 (8 male, 8 female) offspring should be included in each group to achieve α=0.05 with 95% power (G*Power v3.1.9.2). Third, we chose to collect *ex vivo* MR images, as opposed to *in vivo* MR images. The enhanced image quality available with *ex vivo* data increases the statistical power to detect subtle volume changes when performing cross-sectional comparisons of two groups [13, 41]. Therefore, if the study objective is detection of a neuroanatomical phenotype, without attention to its time course, then *ex vivo* imaging is preferable, hence our decision to opt for this approach [13]. Balanced against this argument is the fact that the sample preparation (perfusion and tissue fixation) for *ex vivo* imaging may cause morphological disruption to the tissues, which could affect interpretation of the data. This is particularly true for the ventricular system, which may collapse post-perfusion, such that group-level differences in ventricular volume *in vivo* are not preserved *ex vivo* [45]. This is relevant as *in vivo* studies in MIA rat models do show differences in ventricular volume [20, 24]. Total brain volume and that of most grey matter structures also shrinks post-perfusion [16, 41, 46]. Prior work however, including our own, suggests that major group-level differences in grey matter volumes are preserved despite this shrinkage from *in vivo* to *ex vivo* and can be confirmed *post-mortem* [16, 41]. Taken together, the choice of *ex vivo* MR imaging is consistent with the aims of the current study, but the case for longitudinal *in vivo* studies is also reinforced.

Accepting these limitations, a cautious interpretation of these preliminary data suggests some interesting observations. We found no significant effects of maternal POL exposure on the volume (or microstructure) of the rat hippocampus, prefrontal cortex, corpus striatum or lateral ventricles at PD21. This may suggest that MRI-detectable changes in these regions only emerge with increasing post-natal age and are not present in the juvenile brain [20, 24]. Our preliminary findings highlighting the DD as a structure specifically affected in the juvenile rat brain following MIA are potentially intriguing, based on other lines of evidence. Efferent projections from the DD innervate several brain regions, but the majority project to the hippocampus via the medial septum, where they contribute to the modulation of hippocampal theta (θ) oscillations that are important for attention, spatial and working memory and sensory information processing [47–49]. It is noteworthy then that decreased θ rhythms are reported in the adult rat hippocampus following exposure to MIA at GD15, which are related to memory and sensory processing impairments in these animals [31, 50, 51]. Furthermore, increased number and activity of cholinergic neurons is reported in the basal forebrain (which includes the DD), in the offspring of poly (I:C) exposed dams in early post-natal life (PD1) [52]. Similar findings of increased cholinergic neuron density and activity have also been reported in *post-mortem* human basal forebrain tissue from individuals aged <13 years old with a diagnosis of ASD [53]. Taken together, there is circumstantial evidence to suggest that our preliminary MRI finding of elevated DD volume could have functional and clinical relevance, which can be tested in future studies.

## 5. Conclusion

The findings of the current study provide new insights into the impact of gestational poly (I:C) exposure on the juvenile (PD21) Wistar rat brain, complementing and extending our work using this model [27]. The data, albeit preliminary due to the small sample size, suggests that the DD is the only structure significantly affected at this time-point, with increased volume in offspring exposed to MIA. Our data also provide publicly available data on effect sizes to aid the design of longitudinal *in vivo* MRI studies in this model. Such studies are now clearly required to confirm the functional relevance of the observed DD volume changes with regard to adult behavioural dysfunction and to establish the underlying cellular and molecular mechanisms.

## Author contribution statement

ACV, JCN, and EPP conceived and designed the study with input from MEE and MKH. All animal work, including MIA, was carried out at the University of Manchester by MEE, supervised by MKH, JCN and EPP. TCW optimised the MR pulse sequences used in this study for high-resolution ex vivo MR image acquisition, which was carried out at King’s College London (KCL) by ACV. TCW and ACV carried out the computational and statistical analyses, respectively, of the MR images and atlas-based segmentation data at KCL. ACV drafted and revised the MS with input and the approval of all authors. ACV, JCN and EPP provided financial support for the study.

## Data availability statement

Data collected from this study including the ex vivo MR images may be made freely available upon reasonable requests to the corresponding author.

## Acknowledgements

The authors would like to thank the British Heart Foundation and Dr Po-Wah So (KCL) for supporting and managing the 7T MRI scanner at the King’s College Preclinical Imaging Unit (KCLPIU).

